# Distinctiveness of genes contributing to growth of *Pseudomonas syringae* in diverse host plant species

**DOI:** 10.1101/2020.07.28.216440

**Authors:** Tyler C. Helmann, Adam M. Deutschbauer, Steven E. Lindow

**Author notes:** Emerging Pests and Pathogens Research Unit, Robert W. Holley Center, Agricultural Research Service, United States Department of Agriculture, Ithaca, New York, United States of America. Corresponding author, (SEL).

## Abstract

A variety of traits are necessary for bacterial colonization of the interior of plant hosts, including well-studied virulence effectors as well as other phenotypes contributing to bacterial growth and survival within the apoplast. High-throughput methods such as transposon sequencing (TnSeq) are powerful tools to identify such genes in bacterial pathogens. However, there is little information as to the distinctiveness of traits required for bacterial colonization of different hosts. Here, we utilize randomly barcoded TnSeq (RB-TnSeq) to identify the genes that contribute to the ability of *Pseudomonas syringae* strain B728a to grow within common bean (*Phaseolus vulgaris*), lima bean (*Phaseolus lunatus*), and pepper (*Capsicum annuum*); species representing two different plant families. The magnitude of contribution of most genes to apoplastic fitness in each of the plant hosts was similar. However, 50 genes significantly differed in their fitness contributions to growth within these species. These genes encoded proteins in various functional categories including polysaccharide synthesis and transport, amino acid metabolism and transport, cofactor metabolism, and phytotoxin synthesis and transport. Six genes that encoded unannotated, hypothetical proteins also contributed differentially to growth in these hosts. The genetic repertoire of a relatively promiscuous pathogen such as *P. syringae* may thus be shaped, at least in part, by the conditional contribution of some fitness determinants.

## Introduction

*Pseudomonas syringae* is a ubiquitous species complex containing strains commonly found in association with a variety of both healthy and diseased plants. More than 60 pathovars have been identified in this species, where taxonomic placement is primarily associated with their host range on different groups of host plants [1]. The life cycle of *P. syringae* is typically considered to consist of two overlapping phases: an initial period of epiphytic growth on the surface of plants followed by subsequent invasion and growth in the intracellular spaces (apoplast) [2]. Under favorable environmental conditions, such as either high humidity or surface moisture and moderate temperatures, large epiphytic bacterial populations can establish on leaves, increasing the likelihood that at least some cells will invade the interior of the leaf [3]. After invasion of the leaf through stomata or openings caused by wounds, rapid bacterial multiplication can occur in plants that are susceptible to infection, resulting in the formation of visible leaf symptoms and eventually localized host cell death [2,3]. Most attention to traits involved in plant colonization has focused on those contributing to apoplastic growth. Several such traits including the type III secretion system and certain effector proteins, phytotoxins, siderophores, adhesins, and genes contributing to stress tolerance have been identified [4]. While many genes contribute to growth in particular environments, canonical virulence factors such as type III effectors that contribute to the suppression of the host immune system and establishment of an aqueous environment, as well as phytotoxins, are specifically important to growth in the apoplast [5,6].

Type III effectors in *P. syringae* and other bacterial pathogens are likely to be important determinants of host specificity since they are usually involved in suppression of plant defenses against those strains that are recognized by the plant. The linkage between type III effector repertoires and host range however remains unclear [7]. In addition, individual type III effector genes can be disrupted with little to no effect on virulence, apparently because most of the many effectors produced by a given bacterium have redundant functions [8]. On the other hand, toxins produced by *P. syringae* show little host specificity and typically contribute little to its multiplication in plants [9]. Such toxins instead incite characteristic symptoms that follow infection [9]. While the contribution of such traits to the colonization of a given host plant species has been widely documented, most such studies have been limited to a given pathosystem, and we lack an understanding to what extent such traits contribute to bacterial growth in a plant species-dependent manner.

While the identification of fitness factors of plant pathogenic bacteria by classical gene-by-gene disruption studies is a very laborious process, transposon sequencing (TnSeq), whereby a large mixture of insertional mutants is assayed simultaneously, can quantify the contribution of nearly all nonessential genes to growth in a given plant host. We currently lack an understanding of the extent to which genes that contribute to virulence in one such plant species would also contribute to virulence in other potential host plants. Given that many plant pathogenic bacteria such as *P. syringae* are relatively promiscuous, growing both on and within a variety of often unrelated plant species [10], they might have evolved and retained virulence genes that are selectively important only in a subset of the plants with which they interact. This important question in the ecology and epidemiology of such species can only be answered by determining the extent to which those traits needed for colonization are distinctive to a given host plant. The global assessment of the contribution of genes of a strain to its success in a myriad of different hosts or habitats was however, until recently, challenging.

Random barcoded TnSeq (RB-TnSeq) is a modification of TnSeq in which 20-nucleotide barcodes are associated with distinct individual transposon insertion sites [11], facilitating the repeated interrogation of the same large collection of random insertional mutants for their contribution to fitness in multiple experiments. Because the location of the transposons in the mutant population only needs to be mapped a single time using TnSeq, the composition of the individual members of the mutagenized community can be easily quantified in each of many subsequent experiments by enumeration of only the amplicon barcode regions (BarSeq) [11]. This technique has been used to identify *P. simiae* genes that enable colonization of *Arabidopsis thaliana* roots [12] as well as genes that contributed to competitive growth of *P. syringae* B728a on both the leaf surface and in the apoplast of common bean (*Phaseolus vulgaris*) [13]. Combined with *in vitro* profiling to determine the function of these genes, these high-throughput transposon mutant studies provided considerable insight into the various traits required for plant colonization. A variety of non-essential genes for growth in the bean apoplast including those involved in phytotoxin biosynthesis, the type III secretion system, and alginate biosynthesis were important in growth in the apoplast. In addition to common bean, strain B728a is capable of both multiplication and elicitation of disease symptoms in plant hosts including lima bean (*P. lunatus*) and pepper (*Capsicum annuum*) [10]. Since apoplastic colonization requires growth of the bacteria while in intimate contact with host mesophyll cells, we hypothesized that the genes required to colonize these phylogenetically distinct plant species would differ. Because of the distinct metabolites expected within these plants and the likelihood of differential chemical or physical responses to infection, we hypothesized that a non-overlapping set of genes in *P. syringae* would contribute to its success in these various hosts. Furthermore, we hypothesized that there would be a greater overlap among those genes that contribute most to growth in common bean and lima bean than to those in pepper due to the close phylogenetic relationship of these two *Phaseolus* species in the Fabaceae family compared to that of *C. annuum,* a member of the Solanaceae family, Here, we examine the differential contribution of non-essential genes in *P. syringae* strain B728a to its growth in the apoplast of these three plant species.

## Results

### *P. syringae* B728a growth is supported in diverse plant hosts

Given that the sensitivity of gene fitness measurements of bacteria interacting with plants using methods such as TnSeq is proportional to the extent of multiplication of the bacteria in the plant [13], we measured the growth of *P. syringae* within several possible host plants to determine whether such a strategy could be employed. Population sizes increased approximately 1,000-fold within six days after inoculation in common bean, and over 10,000-fold in both lima bean and pepper (Fig. S1). Such increases indicated that, on average, each inoculated cell had undergone 10 generations of growth in common bean and 13 or more generations of growth in both lima bean and pepper. Lima bean appeared to be relatively more susceptible to infection with *P. syringae* than common bean since it both enabled more extensive growth of the bacterium within its tissues and exhibited more visible disease symptoms (data not shown). In contrast, although *P. syringae* multiplied extensively in pepper, symptoms were restricted to the appearance of only yellow mottling that occurred a week or more after inoculation. In all hosts, infiltration of leaves with bacterial suspensions of 10^5^ cells/ml conferred sufficiently large initial populations of the pathogen (ca. 5 x 10^3^ cells/cm^2^ of leaf) to yield aggregate populations of ca. 10^6^ cells in the collection of several hundred leaves from which cells were subsequently recovered, thus reducing any bottleneck effects. Such initial population sizes were, however, sufficiently low to enable extensive subsequent growth of the inoculated bacterial cells before their recovery from the apoplast.

### Inoculation of a barcoded transposon library in multiple plant hosts

A randomly-barcoded *mariner* transposon library of *P. syringae* B728a described previously [13,14] was inoculated by vacuum infiltration into the leaves of the three plant species to identify genes contributing to host colonization. This mutant library contains 281,417 total strains having insertions mapped in the B728a genome. A subset of 169,826 genic strains in which transposon insertions were in the center 10 – 90% portion of a given gene, were used for calculating gene fitness values. Fitness contributions could be estimated for 4,296 (84%) of the 5,137 protein-coding genes of this strain. Since the mutant library was cultured in King’s medium B (KB) from a frozen stock immediately prior to plant inoculations, any changes in the proportional representation of transposon mutants in a given gene during overnight growth in this condition were used as the control against which growth of the mutants *in planta* was compared.

Samples of the transposon library recovered from each plant host after growth in the plant for 6 days contained at least 68% the total unique barcodes that were present in the inoculum infiltrated into leaves (Table S1). This further supports the results of population sizes after inoculation that indicated that a sufficiently large proportion of the transposon mutant population was successfully introduced into plants during inoculation into the apoplast to avoid a large bottleneck. The contributions of genes to fitness in plants were largely distinct from their contribution to growth in a rich culture medium. Replicate measures of gene fitness values in a given plant host cluster together and separately from those when grown in KB (Fig. 1). It is noteworthy that the patterns of gene fitness values obtained in pepper are more disparate from those seen in both lima bean and common bean. Despite the differential overall growth in these hosts, fitness values ranged from approximately −5.5 to +1.0 in all replicate studies of these 3 host plants (Fig. S2). For each gene, fitness values for the two replicate growth experiments in KB and the three replicate apoplastic growth experiments performed in each plant host were averaged. While there was variation in the standard deviation of the fitness values estimated for a given gene in a particular host, very few genes exhibited high variation in estimated fitness values and the mean standard deviation did not differ between the three plant species tested here (Fig. S3). It is noteworthy that the standard deviation of estimated fitness values of genes that contributed most strongly to either positively or negatively affecting fitness were generally somewhat higher than those of genes that contributed little to fitness in a given host (Fig. S4), suggesting that the fitness contribution of the traits encoded by these genes was perhaps somewhat context–dependent and thus influenced by idiosyncratic features of particular experiments.

**Figure 1.**
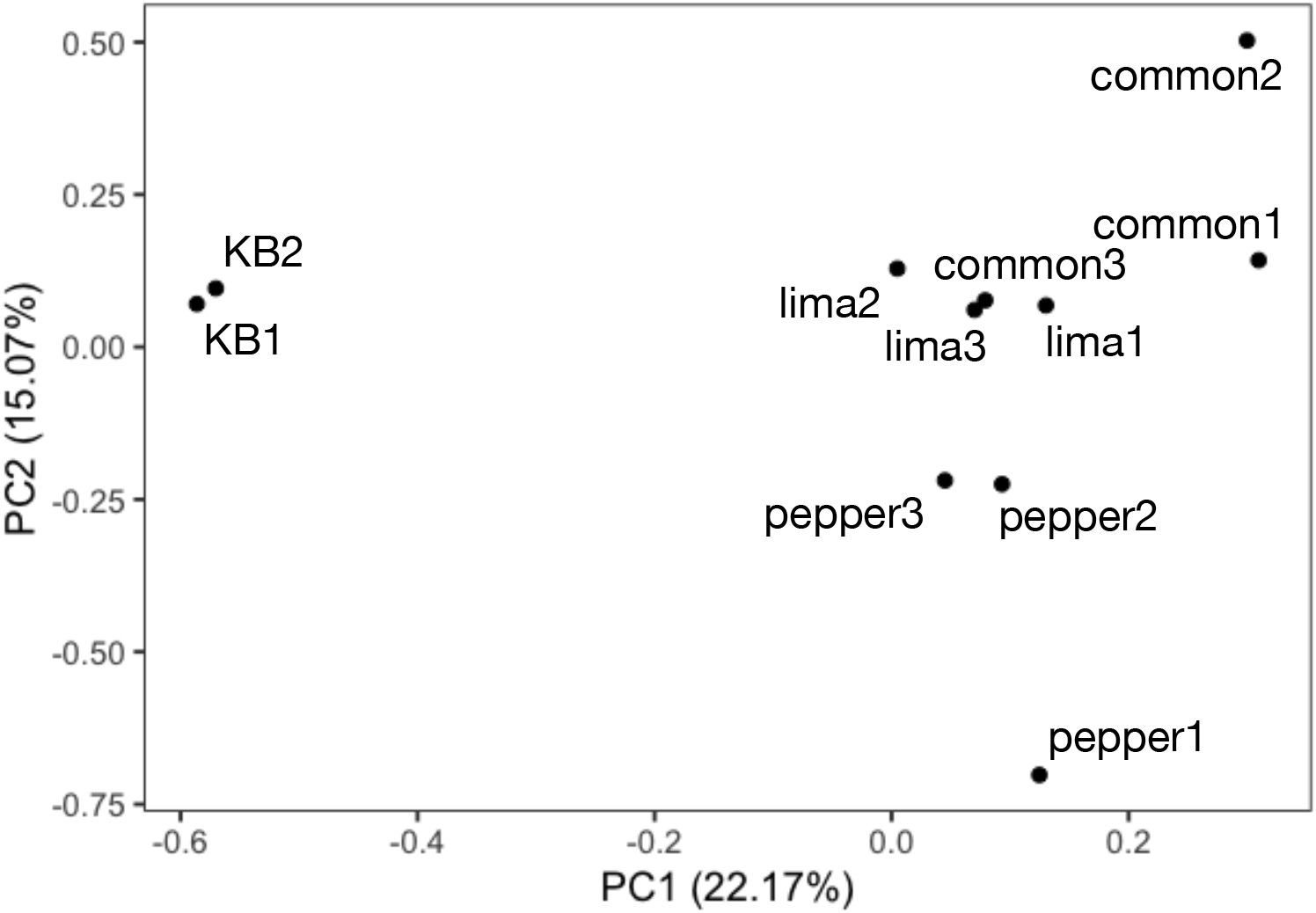
Principle component analysis of *Pseudomonas syringae* B728a grown in King’s B (KB), common bean (“common”), lima bean (“lima”), or pepper, based on gene fitness values calculated for 4,296 genes. Replicate experiments are noted as the numbers 1 to 3.

### Common genes for apoplastic fitness in diverse plant species

To determine the extent to which those genes that had been identified as contributing to apoplastic colonization in common bean [13] were also required for maximum fitness in both lima bean and pepper, we compared gene fitness values after inoculating the mutant library into the apoplast of two additional host plant species. Many of the genes of *P. syringae* that contributed to its fitness in a given host also contributed similarly to its fitness in the other two plant species tested. Pairwise comparisons of gene fitness values on a given plant species with that in other plant species revealed a high correlation between the fitness contribution of a given gene in the apoplast of common bean, lima bean, and pepper (Fig. 2). Pearson correlation coefficients, comparing average fitness values from all genes, ranged from 0.725 (lima bean compared to pepper) to 0.796 (common bean compared to lima bean). While many genes differentially contribute to growth in the apoplast of various plant species, it is clear that nearly as many make similar contributions to fitness on all of the plant species. For both those genes contributing strongly to apoplastic fitness (fitness value < −2) (Fig 3A) or contributing somewhat less to fitness (fitness value < −1) (Fig. 3B) approximately 1/3 contributed to competitive fitness in all three host plants tested. Genes with average fitness values below these two fitness thresholds are listed in Table S2. Those genes with large fitness affects in all hosts are primarily involved in amino acid metabolism, as well as other functional categories including cofactor metabolism, the type III secretion system, and polysaccharide synthesis and regulation (Table 1).

**Figure 2.**
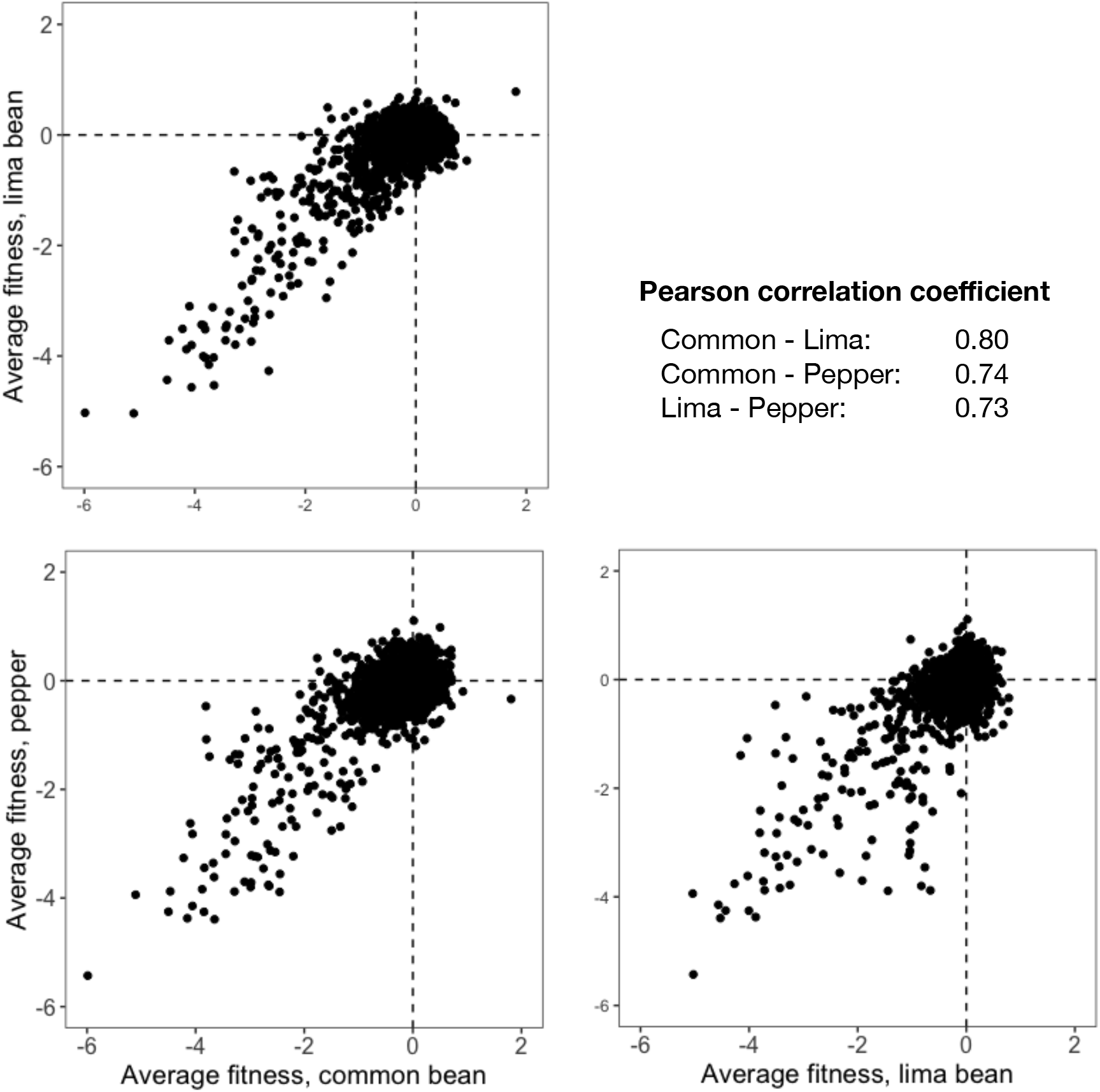
Pairwise comparisons of gene fitness values between plant hosts reveal a high correlation between their contributions to growth of *Pseudomonas syringae* in the apoplast of common bean, lima bean, and pepper.

**Figure 3.**
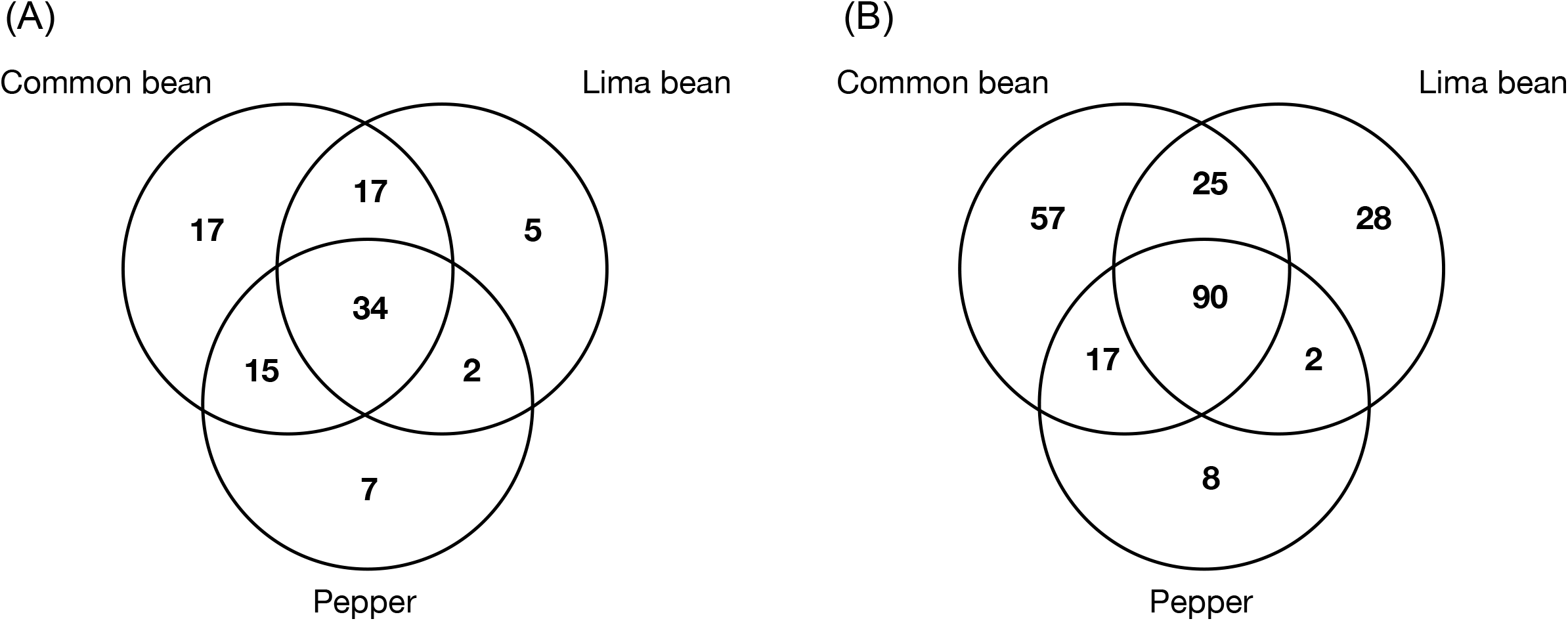
Genes with significant contributions to apoplastic fitness in a given host plant species. Venn diagrams of (A) the number of genes with average fitness values < −2 or (B) the number of genes with average fitness values < −1.

**Table 1.** Number of genes within each functional category with significant contributions to apoplastic fitness in all three host plant species tested. Fitness cutoffs of < −1 and < −2 are both shown. Functional category annotations are primarily based on COG [36] and KEGG [37] annotations, with manual additions and corrections originally published by Yu et al. [33].

| **Functional Category** | **Fitness < -1** | **Fitness < -2** |
| --- | --- | --- |
| Amino acid metabolism and transport | 22 | 15 |
| Unclassified | 14 | 4 |
| Type III secretion system | 12 | 1 |
| Cofactor metabolism | 11 | 0 |
| Polysaccharide synthesis and regulation | 6 | 2 |
| Nucleotide metabolism and transport | 4 | 3 |
| Replication and DNA repair | 4 | 0 |
| Hypothetical | 3 | 2 |
| LPS synthesis and transport | 3 | 2 |
| Carbohydrate metabolism and transport | 2 | 1 |
| Cell division | 2 | 1 |
| Peptidoglycan/cell wall polymers | 2 | 1 |
| Proteases | 2 | 1 |
| Phytotoxin synthesis and transport | 1 | 0 |
| Transcription - Sigma factor | 1 | 1 |
| Transcriptional regulation | 1 | 0 |
| Total | 90 | 34 |

### Differential contributions of genes to apoplastic growth in various plant species

The magnitude of the contribution of 87 genes to apoplastic fitness differed significantly among these 3 plant species (Kruskal-Wallis, p < 0.05). Fifty of those genes had relatively large effects on fitness, having average fitness values of either less than −0.5 or greater than +0.5 in at least one host (Table S3). Those genes that had a differential impact on fitness in different hosts included eight that were annotated as being involved in polysaccharide synthesis and regulation, six in cofactor metabolism, five in phytotoxin synthesis and transport, and four in amino acid metabolism and transport. We also identified six genes that encoded hypothetical proteins that were differentially important in these hosts. The identity of the other 37 genes that contributed differentially to fitness on these hosts, but whose effect was small, having fitness values greater than −0.5 or less than +0.5 in all host plant species are described in Table S4.

### Apparent variations in amino acid availability between plant species revealed by fitness of auxotrophic strains

Genes involved in the biosynthesis of several amino acids were generally required for competitive fitness in the apoplast of all plant species tested (Table 1). However, auxotrophic strains in histidine and isoleucine/valine biosynthesis were less fit in common bean and pepper than in lima bean (Fig. 4) suggesting that these amino acids were more abundant in lima bean than in the other two species. Within these pathways, average fitness values for *hisD, hisH,* and *ilvHI* differed significantly in at least one plant species (Kruskal-Wallis, p < 0.05).

**Figure 4.**
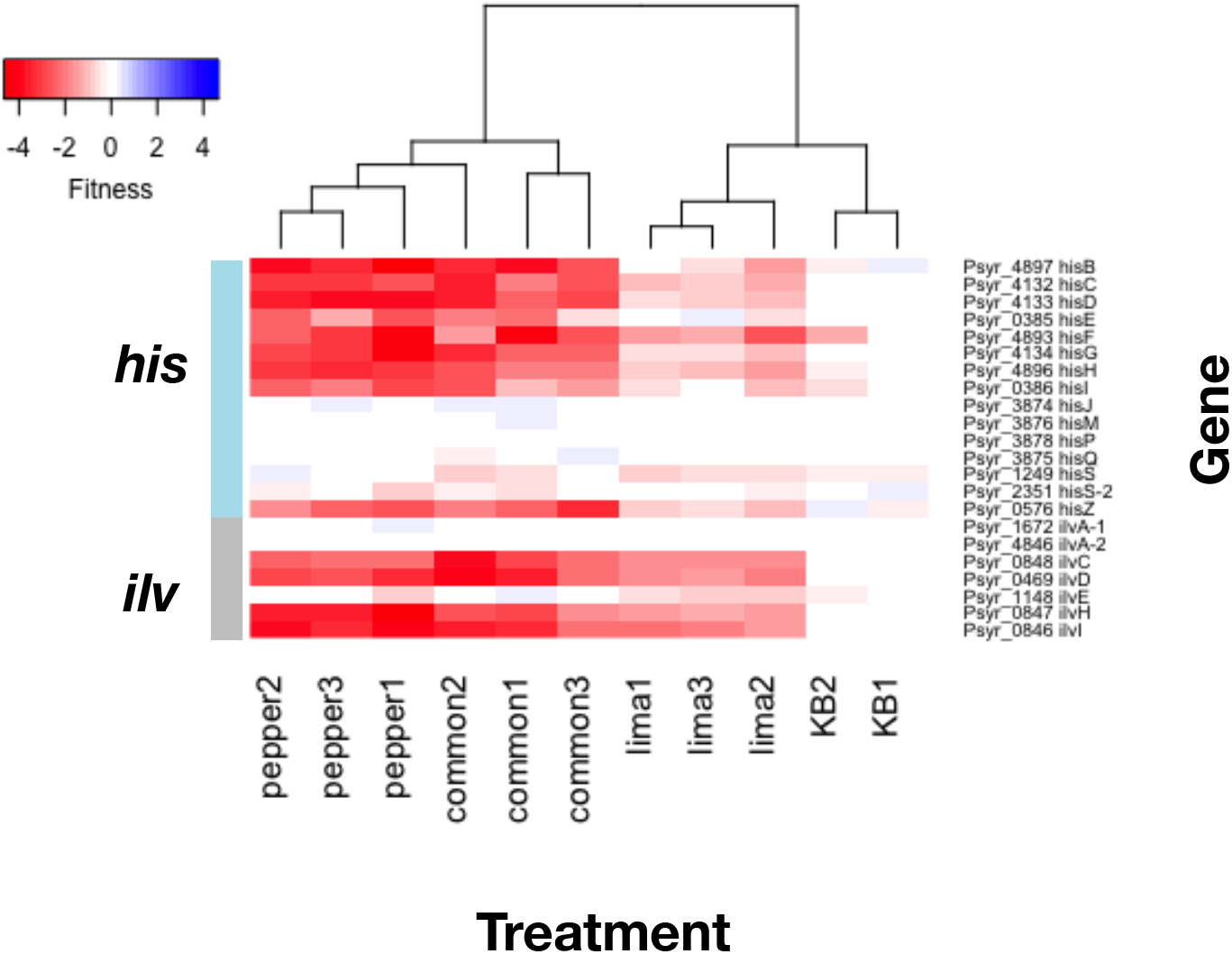
Heatmap of apoplastic fitness contribution of histidine and isoleucine/valine biosynthetic genes to growth in various plant species. Genes *hisD, hisH,* and *ilvHI* had significantly different fitness values in different plant species by a Kruskal-Wallis test (p < 0.05).

### Phytotoxin production appears to be a host-specific virulence trait

While the production of phytotoxins such as syringomycin contribute to competitive fitness in the apoplast of common bean [13], it is noteworthy that such genes contributed less to fitness in lima bean than in both common bean and pepper (Fig. 5) suggesting that this host was less susceptible to the damage caused by phytotoxins such as syringomycin. The average fitness values of genes encoding the regulatory proteins SalA, SylA, and SyrP, as well as those encoding the secretion protein PseB and the amino acid adenylation protein SypA differed significantly among these three plant species (Kruskal-Wallis p < 0.05).

**Figure 5.**
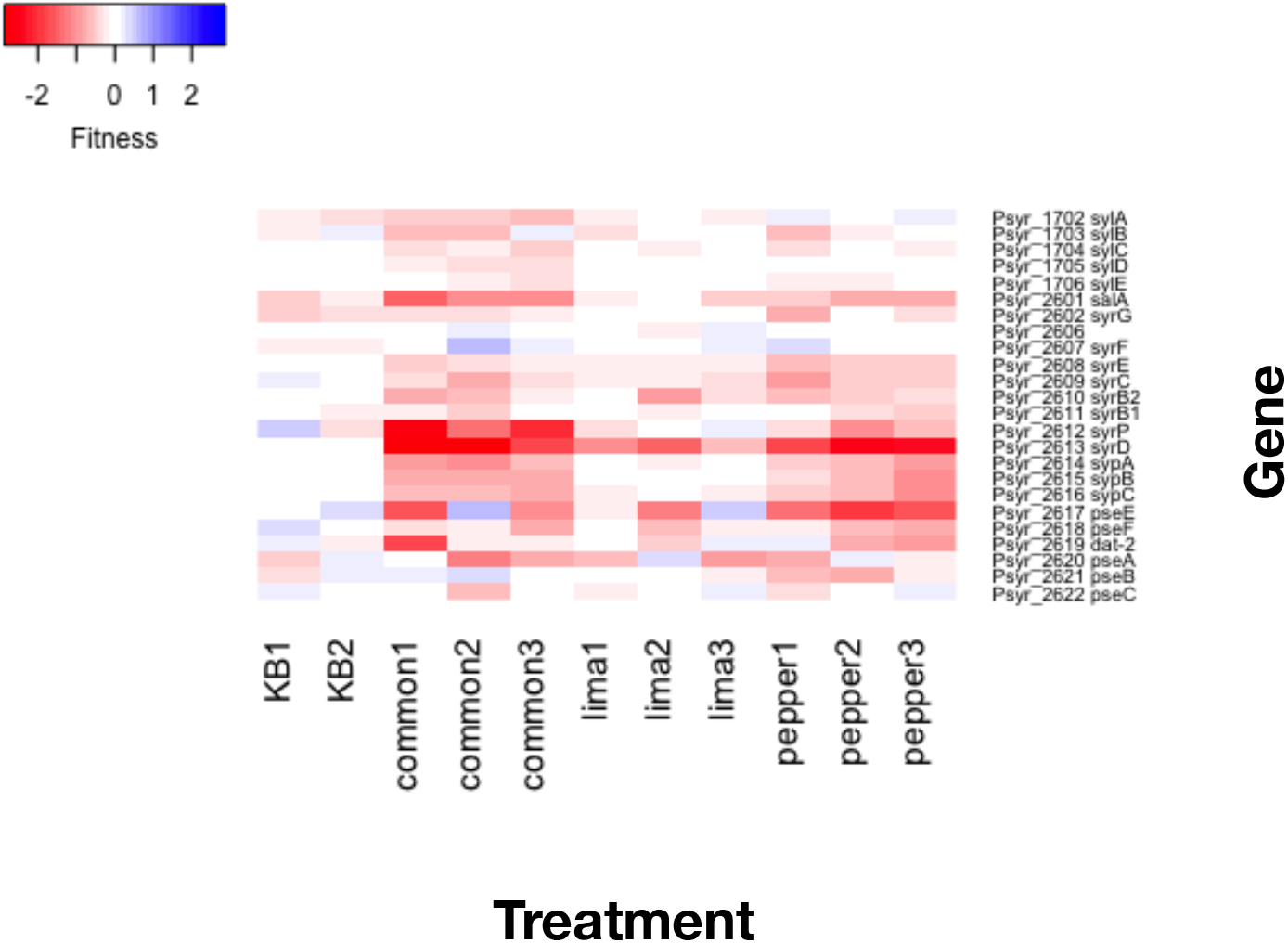
Heatmap of apoplastic fitness values of genes involved in phytotoxin biosynthesis. Genes *sylA, salA, syrP, sypA,* and *pseB* had significantly different fitness values in these 3 plant species by a Kruskal-Wallis test (p < 0.05).

### Alginate production in *P. syringae* varies in importance among host plants

While polysaccharide biosynthesis and regulation is considered a broadly important plant colonization trait, the genes involved in the biosynthesis of alginate contributed differentially to apoplastic growth in the three plant species investigated (Fig. 6). Specifically, the alginate biosynthesis genes *alg44, alg8, algA-1, algF, algI,* and *algK* were more important in the colonization of lima bean and common bean than in pepper (Kruskal-Wallis, p < 0.05).

**Figure 6.**
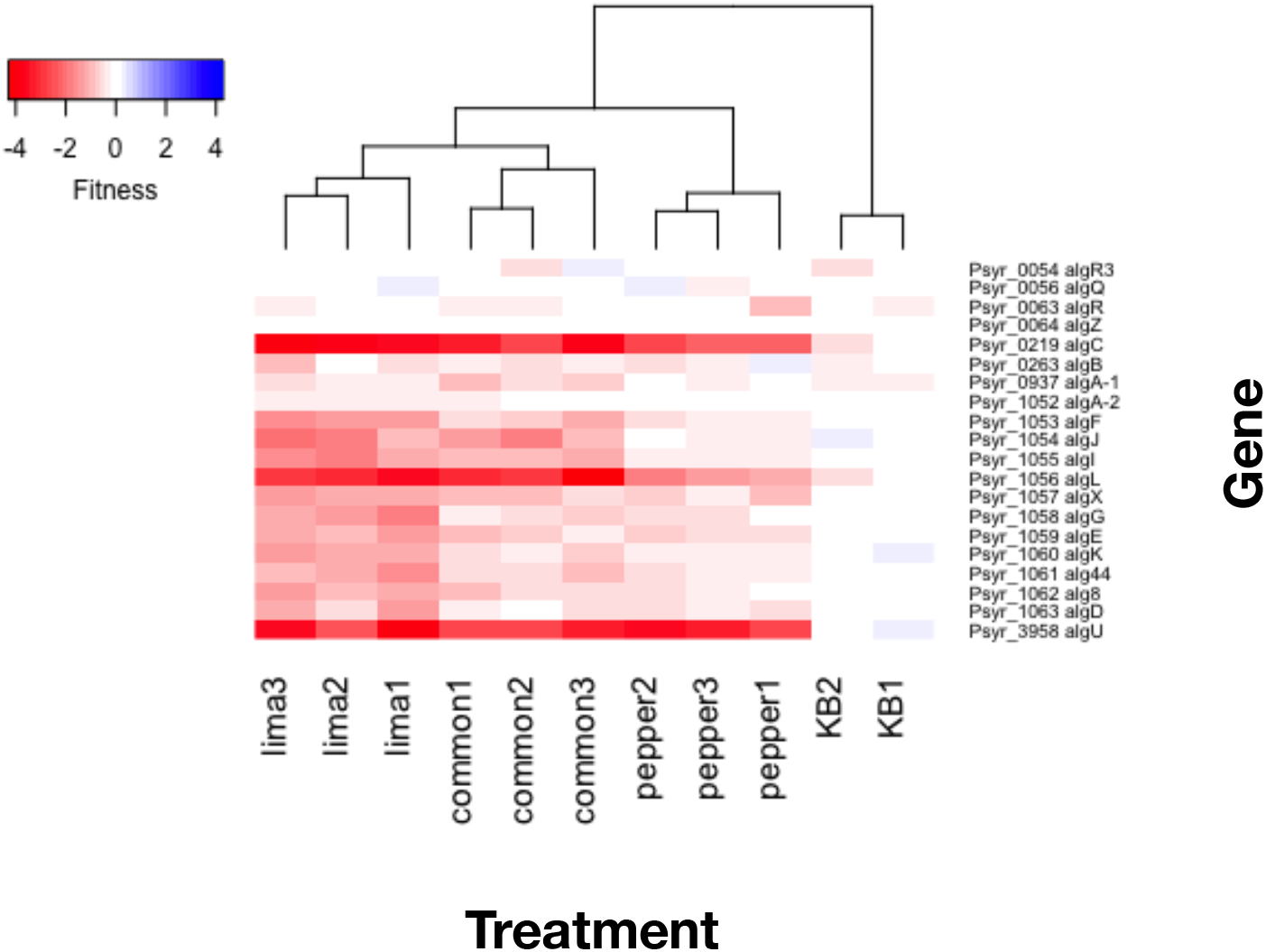
Heatmap of apoplastic fitness for alginate biosynthetic genes. Genes *alg44, alg8, algA-1, algF, algI,* and *algK* had significantly different fitness values in the 3 plant species by a Kruskal-Wallis test (p < 0.05).

## Discussion

RB-TnSeq is a powerful method that has been used in a variety of bacterial species to identify those genes that contribute to fitness in diverse *in vitro* conditions [11,14,15]. In one TnSeq study, about 100 genes were found to either increase or decrease the fitness of *Dickeya dadantii* in chicory [16], highlighting the importance of the biosynthetic pathways for leucine, cysteine, and lysine for bacterial growth within the plant. A similar study in *Pantoea stewartii* ssp. *stewartii* identified genes that were important for its growth within corn (*Zea mays*) xylem [17]. Typical of most studies, these reports focused on a single pathosystem. This limited use of this powerful technology is largely due to the cost and effort required to map genomic insertions in the mutagenized population at the start and end of each setting to which the mutagenized population is exposed. In contrast, after initially associating several uniquely tagged transposons with a given gene in RB-TnSeq the relative proportions of the different insertional mutants can be readily assessed by quantifying the amplified barcodes. In this study, interrogation of a mutant mixture after growth in three different host plants provided confident estimates of the differential contribution of each non-essential gene to fitness in these hosts. This proved to be a very powerful tool to generate hypotheses regarding host-specific virulence traits by identifying both common and host-specific genes that enable growth in the intercellular spaces of plants.

We hypothesized that at least a subset of the genes in *P. syringae* required for colonization of the apoplast of a given host plant would not be required in other plants, with the assumption that distinctive features of the niche in the interior of these plants would differ in resources and chemical defenses. A larger number of genes are required for colonization of the apoplast of common bean compared to the leaf surface [13], suggesting that the leaf interior provides a more complex chemical or physical habitat than the leaf surface. Through fitness profiling of amino acid auxotrophs, the importance of various amino acid biosynthetic pathways for growth both on the leaf surface and in the apoplast of common bean has been demonstrated. This suggested that some amino acids are less abundant than others in both of these environments, supporting direct measurements made by others [18]. We find here that the relative abundance of available amino acids in the apoplast seems to also vary between plant species. Specifically, the intercellular spaces of lima bean appears to contain more accessible histidine, isoleucine, and valine than in both common bean and pepper since genes for the production of these resources played a lesser role in *P. syringae* than in the other plant species (Fig. 4). On the other hand, similar relative amounts of other amino acids are apparently present in the three plant species tested, as the genes in other amino acid biosynthetic pathways did not contribute to fitness differentially in plants examined here.

While they are not essential for growth *in planta, P. syringae* produces various phytotoxins that contribute to symptom development in plants [19,20]. Strain B728a produces the lipodesinonapeptides syringomycin and syringopeptin, as well as the peptide derivative syringolin A [21,22]. Syringolin A counteracts stomatal immunity through proteasome inhibition [23], and genes in this biosynthetic pathway did not contribute to apoplastic growth in any of the three plant studied here. Syringomycin and syringopeptin function as pore-forming cytotoxins, contributing to ion leakage [24]. It was interesting to note that the biosynthetic genes for these non-specific toxins contributed more to fitness in common bean and pepper than in lima bean (Fig. 5). Curiously, while strain B728a grew to higher population sizes in pepper than in common bean (Fig. 1), symptom development is much less apparent on pepper than in either common or lima bean, without the appearance of any necrosis. While these three plant species were all susceptible to disease caused by strain B728a, lima bean appeared much more susceptible than common bean because of the substantially higher apoplastic populations that were attained as well as an earlier onset and more extensive necrotic lesion formation that it supported (Fig. S1). Thus, while it appears that syringomycin is a general virulence trait required for maximum growth within plants, and not merely a factor contributing to necrotic lesion development, the mechanism by which it contributes to growth in plants remains unclear. It is possible that differences in the immune responses between these plant species might be modulated by syringomycin, thus contributing to the differential contribution of this trait to aplastic growth.

Given that many genes contributed similarly to virulence in all three hosts, it remains unclear why *P. syringae* strain B728a multiplied to substantially different population sizes in these plants. Low water availability on both the leaf surface and in the apoplast limits bacterial growth [25]. In response, *P. syringae* apparently upregulates the production of the exopolysaccharide alginate, to help maintain available moisture for bacterial growth [1]. Alginate production is an important factor for both virulence and epiphytic fitness of *P. syringae* [1,26]. While alginate biosynthetic mutants were less fit than most other mutants in the apoplast of all three hosts tested in this study, alginate appears to be a more important virulence trait in lima bean relative to that in pepper. In *P. syringae* pv. *tomato* DC3000, alginate biosynthetic genes are expressed in both host and non-host plants [27]. While alginate production is apparently a generally important virulence trait, differences in its relative contribution to virulence in these plant species may indicate variation in the relative moisture content in the apoplastic space of lima bean and pepper. It is possible that the anatomical differences between these plants contribute to differences in water availability. For example, pepper leaves are thicker than that of either of the two bean species, and its water loss during the infection process may be somewhat less than in lima bean, thus enabling its leaves to better retain water that may be released by the plant to the apoplast during the infection process, diminishing a requirement for alginate production by *P. syringae* to maintain hydration [6].

Given that *P. syringae* strain B728a is capable of symptom formation in several additional plant species beyond those tested here [10], it should prove informative to interrogate the contribution of both the common and unique virulence genes identified here in a more disparate collection of plant species that presumably would provide yet more distinctive plant habitats that would require additional host-specific fitness traits than those revealed here. Further studies of interactions with the plant species examined here should also enable the identification of additional genes that play a quantitatively smaller, but ecologically important role in their colonization. The fitness values observed for many genes reflected 50% or less growth of the disruption mutants compared to the wild-type strain. Such a large effect of these genes on growth over the relatively few (10 to 13 generations) in these studies suggests that there is strong selection for these genes in plants, especially when considering the many generations that the species would undergo in a given year. Because of the limited number replications used in the study we did not use a multiple testing correction for the Kruskal-Wallis tests because the resulting p-value (10^-5^) was deemed to be too conservative. This risks the identification of some genes as “host-specific” when they were not. However, it is important to note that all of the genes that contributed differentially to colonization of these three plant species (Table 2) were found to be required for fitness in at least one of these plant species, having an average fitness value less than −0.5 in at least one host. There is less support for a few other genes for which differential contribution fitness was indicated (Table 3). The contribution of such genes with lesser effects on virulence might be better quantified in further studies in which the mutant mixtures are recovered after infection events and used to re-inoculate additional plants, thereby increasing the number of generations of multiplication over which fitness values could be estimated.

*P. syringae* is a commonly observed phyllosphere resident that is also found in other location in the global water cycle [28]. As such, it presumably interacts with a large number of different plants, both as a surface colonist and after entering into plants [29]. These various habitats presumably require very different genes to enable it to exploit these diverse settings. There is therefore much to learn about habitat-specific contributions of its many genes. The high-throughput analysis of insertional mutant mixtures used here not only reveals those traits important in a given setting, but also elucidates the limiting factors in those habitats. As such, further exploitation of RB-TnSeq enables functional genomics to be applied to *P. syringae* in its diverse natural habitats. While we considered here the contribution of genes to fitness in several of the many plants in which *P. syringae* might find itself, there are many other settings in which a subset of its genes may become particularly important to fitness. Clearly, *P. syringae* has maintained genes preferentially important in some hosts, and many other genes may play similar habitat-specific roles.

## Materials and Methods

### Bacterial strains and growth media

*P. syringae* pv. *syringae* B728a was originally isolated from a bean leaf (*Phaseolus vulgaris*) in Wisconsin [30]. The complete genome for B728a is available on NCBI GenBank as accession CP000075.1 [21]. B728a was grown on King’s B (KB) agar or in KB broth [31], at 28°C. When appropriate, the following antibiotics were used at the indicated concentrations: 100 μg/ml rifampicin, 100 μg/ml kanamycin, and 21.6 μg/ml natamycin (an anti-fungal). Strains used in this study are listed in Table S5. Primers used for BarSeq amplification are described in [11].

### Plant growth conditions

Common bean (*P. vulgaris* var. Blue Lake Bush 274) and lima bean (*P. lunatus* var. Haskell) were grown in Super Soil (Scotts Miracle-Gro), at a density of five to seven plants per 10 cm diameter pot, in a greenhouse for two weeks before inoculation. Pepper (*C. annuum* var. Cal Wonder) were grown in 10 cm diameter pots (three to five plants per pot) containing Super Soil and grown in a greenhouse for ca. 6 weeks to a height of ca. 30 cm. Leaves were kept dry to minimize epiphytic bacterial populations. Metal halide lights (1000 W) were used to provide supplemental lighting for a 16-hour day length.

### Bacterial apoplastic growth measurements

Wild type strain B728a was grown overnight on KB with rifampicin, washed in 10 mM KPO_4_ (pH 7.0), and adjusted to a cell concentration of 2×10^5^ CFU/ml in 1 mM KPO4. Cells were inoculated into leaves using a blunt syringe. Leaf samples (3 discs per leaf) were excised using a 5 mm-diameter cork borer and placed into microfuge tubes containing 200 μl 10 mM KPO_4_ and two 3 mm glass beads, and macerated by shaking for 30 seconds at 2400 rpm in a Mini-Beadbeater-96 (Biospec Products) before dilution plating of appropriate serial dilutions on KB containing rifampicin and natamycin. Colonies were enumerated following two days growth at 28°C.

### Library recovery and growth in KB

For each inoculation, a 1.25 ml aliquot of a glycerol stock containing the transposon library that had been stored at −80°C was placed in 25 ml fresh KB with kanamycin and grown for approximately 7 hours at 28°C with shaking until the culture reached mid-log phase (OD_600_ 0.5 - 0.7). Time0 samples enumerating the relative abundance of each mutant were collected from the inoculum; 1 ml aliquots were pelleted by centrifugation and the pellets frozen until DNA purification. The remaining cells were then washed twice in 10 mM KPO_4_ prior to plant inoculation.

To determine the growth of the mutant library in KB, a 50 μl log phase cell culture (OD_600_ 0.5) was inoculated into 950 μl KB with kanamycin in a 24-well plate. The plate was incubated at 28°C with shaking for 15 hours. Cells were collected by centrifugation, and frozen prior to DNA purification. Calculation of the fitness values from the KB control treatment is discussed in [13].

### Inoculations of the transposon library into plants

Washed cells were re-suspended to a concentration of 2×10^5^ CFU/ml in 1 mM KPO_4_. The soil of potted plants was covered with cotton to hold the soil in place, and the foliage was inverted into open containers containing ca. 1.5 L of bacterial cell suspension in an open glass bell jar. A vacuum was then applied to the plants sealed within the bell jar for 1.25 minutes and then removed rapidly to force the inoculum into the evacuated apoplast. Ca. 100 pots of plants of a given species were inoculated for a given replicate experiment. Plants were allowed to dry overnight and then moved to the greenhouse for six days.

### Bacterial isolation from the apoplast

Leaves of each plant species were excised from the plants and then chopped in a blender to yield fragments with an average diameter of about 1 to 4 mm. The slurry of leaf fragments was then placed in a water-filled glass dish and placed in a sonication water bath (Branson 5510, output frequency 40 kHz) for 15 min to remove cells. The resulting slurry-containing cells that were dislodged from the leaf interior was filtered through a coffee filter to remove most plant debris. 10% of the ca. 5-10 L of buffer containing the largely plant-free cell suspension was then subjected to sequential additional filtration steps of Whatman filters (20 μm, 10 μm, and 6 μm) to remove much of the remaining plant debris. Bacterial cells were then removed from suspension by centrifugation at 4696 x g for 10 minutes. The pellet was re-suspended in water, and aliquots of cell pellets were frozen prior to DNA purification.

### DNA isolation and library preparation

DNA from frozen pellets was isolated using the Qiagen DNeasy Blood & Tissue Kit according to manufacturer’s instructions. Cell lysis was done at 50°C for 10 minutes as per optional instructions. For those samples having excess residual plant material, lysed cells were centrifuged at 1,500 x g for 5 minutes before loading the supernatant onto purification columns. Purified genomic DNA was measured on a nanodrop device and 200 ng of total DNA was used as a template for DNA barcode amplification and adapter ligation as established previously [11]. For each time0 and plant experimental sample, two separately purified DNA samples were sequenced as technical replicates.

### Sequencing and fitness value estimation

Barcode sequencing, mapping, and analysis to calculate the relative abundance of barcodes was done using the RB-TnSeq methodology and computation pipeline developed by Wetmore *et al.* [11]; code available at bitbucket.org/berkeleylab/feba/. TnSeq was used to map the insertion sites and associate the DNA barcodes to these insertions, as described in [13]. Unique barcoded mutants were identified as those that both mapped to the genome and for which 3 or more reads were obtained during sequencing of the amplified barcodes in a given experiment. For each experiment, fitness values for each gene were calculated as the log_2_ of the ratio of relative barcode abundance following library growth in a given condition divided by relative abundance in the time0 sample. Fitness values were normalized across the genome so the typical gene had a fitness value of 0. Fitness values from sequencing replicates were averaged for each experiment. All experiments passed previously described quality control metrics [11], with the exception of one technical replicate (*P.vulgaris_2*), which was removed from analysis. The requirements for a successful experiment included the detection of more than 50 median reads per gene and a consistency in the calculated fitness value estimated from mutants with insertions in the 5’ and 3’ distal halves of a given gene [11]. Experimental fitness values are publically available at fit.genomics.lbl.gov. Fitness values calculated for growth in common bean were previously generated [13].

### Genomic fitness data analysis

All analysis of gene fitness values and gene metadata was done in R [32]. A PCA plot of experiments was generated from the matrix of gene fitness values using the function prcomp. To better classify genes based on their genomic annotation, we assigned gene names, gene product descriptions, and broad functional categories based on the previously annotated genomic metadata [33]. For each gene, fitness values for experimental replicates were averaged to calculate an average gene fitness value for each plant species. A Kruskal-Wallis test was used to identify those genes in which the median fitness in at least one plant host was significantly different than that in other plant species (p < 0.05). Fitness values in KB were not included in this analysis. From these results, genes were removed to a separate list if the absolute average fitness was less than 0.5 in all conditions. Graphs were plotted in R using the ggplot2 package, version 3.1.1 [34]. Heatmaps were plotted in R using the gplots package, version 3.0.1.1 [35].

## Acknowledgements

We thank Morgan Price for assistance with RB-TnSeq sequence analysis. Plants were grown and maintained with the assistance of the UC Berkeley greenhouse staff, led by Tina Wistrom. Funding for TCH was partially provided by the Arnon Graduate Fellowship and the William Carroll Smith Fellowship. This work used the Vincent J. Coates Genomics Sequencing Laboratory at UC Berkeley, supported by NIH S10 OD018174 Instrumentation Grant.

## References

1. Xin XF, Kvitko BH, He SY. *Pseudomonas syringae:* what it takes to be a pathogen. Nat Rev Microbiol. 2018;16: 316–328. doi:10.1038/nrmicro.2018.17

2. Melotto M, Underwood W, He SY. Role of stomata in plant innate immunity and foliar bacterial diseases. Annu Rev Phytopathol. 2008;46: 101–122. doi:10.1146/annurev.phyto.121107.104959

3. Hirano SS, Upper CD. Bacteria in the leaf ecosystem with emphasis on *Pseudomonas syringae* - a pathogen, ice nucleus, and epiphyte. Microbiol Mol Biol Rev. 2000;64: 624–653. doi:10.1128/mmbr.64.3.624-653.2000

4. Lindeberg M, Myers CR, Collmer A, Schneider DJ. Roadmap to new virulence determinants in *Pseudomonas syringae:* insights from comparative genomics and genome organization. Mol Plant-Microbe Interact. 2008;21: 685–700. doi: 10.1094/mpmi-21-6-0685

5. Nomura K, Melotto M, He SY. Suppression of host defense in compatible plant-*Pseudomonas syringae* interactions. Curr Opin Plant Biol. 2005;8: 361–368. doi:10.1016/j.pbi.2005.05.005

6. Xin XF, Nomura K, Aung K, Velásquez AC, Yao J, Boutrot F, et al. Bacteria establish an aqueous living space in plants crucial for virulence. Nature. 2016;539: 524–529. doi:10.1038/nature20166

7. Alfano JR, Collmer A. Type III secretion system effector proteins: double agents in bacterial disease and plant defense. Annu Rev Phytopathol. 2004;42: 385–414. doi:10.1146/annurev.phyto.42.040103.110731

8. Cunnac S, Lindeberg M, Collmer A. *Pseudomonas syringae* type III secretion system effectors: repertoires in search of functions. Curr Opin Microbiol. 2009;12: 53–60. doi:10.1016/j.mib.2008.12.003

9. Alfano JR, Collmer A. Bacterial pathogens in plants: Life up against the wall. Plant Cell. 1996;8: 1683–1698. doi:10.1105/tpc.8.10.1683

10. Morris CE, Lamichhane JR, Nikolić I, Stanković S, Moury B. The overlapping continuum of host range among strains in the Pseudomonas syringae complex. Phytopathol Res. 2019;1: 1–16. Available: doi:10.1186/s42483-018-0010-6

11. Wetmore KM, Price MN, Waters RJ, Lamson JS, He J, Hoover CA, et al. Rapid quantification of mutant fitness in diverse bacteria by sequencing randomly bar-coded transposons. MBio. 2015;6: 1–15. doi:10.1128/mBio.00306-15

12. Cole BJ, Feltcher ME, Waters RJ, Wetmore KM, Mucyn TS, Ryan EM, et al. Genome-wide identification of bacterial plant colonization genes. PLoS Biol. 2017;15: 1–24. doi:10.1371/journal.pbio.2002860

13. Helmann TC, Deutschbauer AM, Lindow SE. Genome-wide identification of *Pseudomonas syringae* genes required for fitness during colonization of the leaf surface and apoplast. Proc Natl Acad Sci. 2019;116: 18900–18910. doi: 10.1073/pnas.g273cl:3981908858116

14. Helmann TC, Ongsarte CL, Lam J, Deutschbauer AM, Lindow SE. Genome-wide transposon screen of a *Pseudomonas syringae mexB* mutant reveals the substrates of efflux transporters. MBio. 2019;10: e02614–19. doi:10.1128/mBio.02614-19

15. Price MN, Wetmore KM, Waters RJ, Callaghan M, Ray J, Liu H, et al. Mutant phenotypes for thousands of bacterial genes of unknown function. Nature. 2018;557: 503–509. doi:10.1038/s41586-018-0124-0

16. Royet K, Parisot N, Rodrigue A, Gueguen E, Condemine G. Identification by Tn-seq of *Dickeya dadantii* genes required for survival in chicory plants. Mol Plant Pathol. 2019;20: 287–306. doi:10.1111/mpp.12754

17. Duong DA, Jensen R V., Stevens AM. Discovery of *Pantoea stewartii* ssp. *stewartii* genes important for survival in corn xylem through a Tn-Seq analysis. Mol Plant Pathol. 2018;19: 1929–1941. doi:10.1111/mpp.12669

18. O’Leary BM, Neale HC, Geilfus CM, Jackson RW, Arnold DL, Preston GM. Early changes in apoplast composition associated with defence and disease in interactions between *Phaseolus vulgaris* and the halo blight pathogen *Pseudomonas syringae* pv. *phaseolicola*. Plant Cell Environ. 2016;39: 2172–2184. doi:10.1111/pce.12770

19. Bender CL, Alarcon-Chaidez F, Gross DC. *Pseudomonas syringae* phytotoxins: mode of action, regulation, and biosynthesis by peptide and polyketide synthetases. Microbiol Mol Biol Rev. 1999;63: 266–292.doi:10.1117/1.OE.55.11.115102

20. Xu GW, Gross DC. Evaluation of the role of syringomycin in plant pathogenesis by using Tn5 mutants of Pseudomonas syringae pv. syringae defective in syringomycin production. Appl Environ Microbiol. 1988;54: 1345–1353.

21. Feil H, Feil WS, Chain P, Larimer F, DiBartolo G, Copeland A, et al. Comparison of the complete genome sequences of *Pseudomonas syringae* pv. *syringae* B728a and pv. *tomato* DC3000. Proc Natl Acad Sci USA. 2005;102: 11064–11069. doi: 10.1073/pnas.0504930102

22. Grgurina I, Mariotti F, Fogliano V, Gallo M, Scaloni A, Iacobellis NS, et al. A new syringopeptin produced by bean strains of *Pseudomonas syringae* pv. *syringae*. Biochim Biophys Acta - Protein Struct Mol Enzymol. 2002;1597: 81–89. doi: 10.1016/S0167-4838(02)00283-2

23. Schellenberg B, Ramel C, Dudler R. *Pseudomonas syringae* virulence factor syringolin A counteracts stomatal immunity by proteasome inhibition. Mol Plant Microbe Interact. 2010;23: 1287–1293. doi:10.1016/0584-8539(93)80243-4

24. Hutchison ML, Gross DC. Lipopeptide phytotoxins produced by *Pseudomonas syringae* pv. *syringae:* comparison of the biosurfactant and ion channel-forming activities of syringopeptin and syringomycin. Mol Plant-Microbe Interact. 1997;10: 347–354. doi: 10.1094/mpmi.1997.10.3.347

25. Beattie GA. Water relations in the interaction of foliar bacterial pathogens with plants. Annu Rev Phytopathol. 2011;49: 533–555. doi:10.1146/annurev-phyto-073009-114436

26. Yu J, Penaloza-Vázquez A, Chakrabarty AM, Bender CL. Involvement of the exopolysaccharide alginate in the virulence and epiphytic fitness of *Pseudomonas syringae* pv. *syringae*. Mol Microbiol. 1999;33: 712–720. doi:10.1046/j.1365-2958.1999.01516.x

27. Keith RC, Keith LMW, Hernández-Guzmán G, Uppalapati SR, Bender CL. Alginate gene expression by *Pseudomonas syringae* pv. *tomato* DC3000 in host and non-host plants. Microbiology. 2003;149: 1127–1138. doi:10.1099/mic.0.26109-0

28. Morris CE, Monteil CL, Berge O. The life history of *Pseudomonas syringae:* Linking agriculture to Earth system processes. Annu Rev Phytopathol. 2013;51: 85–104. doi: 10.1146/annurev-phyto-082712-102402

29. Wilson M, Hirano SS, Lindow SE. Location and survival of leaf-associated bacteria in relation to pathogenicity and potential for growth within the leaf. Appl Environ Microbiol. 1999;65: 1435–1443.

30. Loper JE, Lindow SE. Lack of evidence for *in situ* fluorescent pigment production by *Pseudomonas syringae* pv. *syringae* on bean leaf surfaces. Phytopathology. 1987;77: 1449–1454. doi:10.1094/Phyto-77-1449

31. King EO, Ward MK, Raney DE. Two simple media for the demonstration of pyocyanin and fluorescin. J Lab Clin Med. 1954;44: 301–307. doi: 10.5555/uri:pii:002221435490222X

32. R Core Team. R: A language and environment for statistical computing. Vienna, Austria: R Foundation for Statistical Computing; 2017. doi:10.1007/978-3-540-74686-7

33. Yu X, Lund SP, Scott RA, Greenwald JW, Records AH, Nettleton D, et al. Transcriptional responses of *Pseudomonas syringae* to growth in epiphytic versus apoplastic leaf sites. Proc Natl Acad Sci. 2013;110: E425–E434. doi: 10.1073/pnas.1221892110

34. Wickham H. ggplot2: Elegant Graphics for Data Analysis. Springer-Verlag New York; 2016. Available: https://ggplot2.tidyverse.org

35. Warnes GR, Bolker B, Bonebakker L, Gentleman R, Huber W, Liaw A, et al. gplots: Various R Programming Tools for Plotting Data. 2019. Available: https://cran.r-project.org/package=gplots

36. Chen IMA, Chu K, Palaniappan K, Pillay M, Ratner A, Huang J, et al. IMG/M v.5.0: an integrated data management and comparative analysis system for microbial genomes and microbiomes. Nucleic Acids Res. 2019;47: D666–D677. doi: 10.1093/nar/gky901

37. Kanehisa M, Goto S. KEGG: Kyoto Encyclopedia of Genes and Genomes. Nucleic Acids Res. 2000;28: 27–30. doi:10.1016/j.meegid.2016.07.022

