## Supplementary Information for "Distinctiveness of genes contributing to growth of *Pseudomonas syringae* in diverse host plant species"

^#a^Current address: USDA, Agricultural Research Service, Robert W. Holley Center, Emerging Pests and Pathogens Research Unit, 538 Tower Road, Ithaca, NY, USA 14853.

* Corresponding author

**ORCID**

Tyler C. Helmann, 0000-0002-8431-6461

Adam M. Deutschbauer, 0000-0003-2728-7622

Steven E. Lindow, 0000-0001-8333-6674

**Supplementary data and figures**

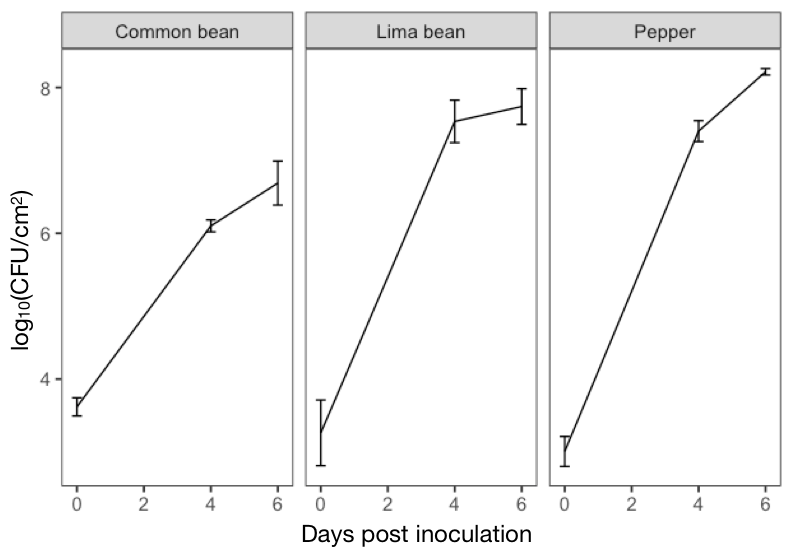

Figure S1. Growth of *P. syringae* B728a in the apoplast of the susceptible hosts common bean (*Phaseolus vulgaris*), lima bean (*P. lunatus*), and pepper (*Capsicum annuum*). The vertical bars represent the standard deviation of the mean.

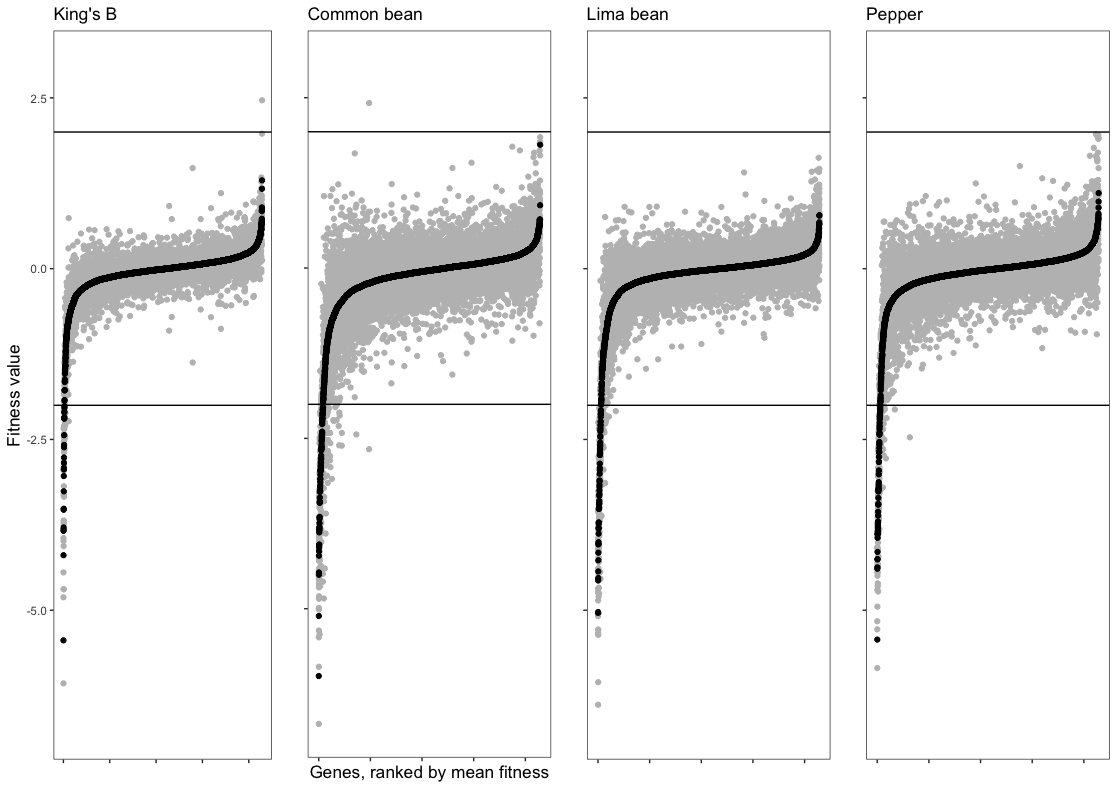

Figure S2. Rank ordered mean gene fitness values for each plant species and the KB control medium in which *P. syringae* was grown. Fitness values for independent replicate experiments are shown in grey, while mean fitness values are shown in black. Gene fitness value is calculated as the log_2_ of the ratio of the barcode counts following growth in a given setting compared to the barcode counts before inoculation. Black lines indicated at fitness values of -2 and +2 are used to reveal strong phenotypes; for example a value of -2 indicates that mutants are 25% as fit as the typical strain in the mutant library. In each dataset, fitness values < -2 or > +2 are more than three standard deviations from the mean (approximately 0).

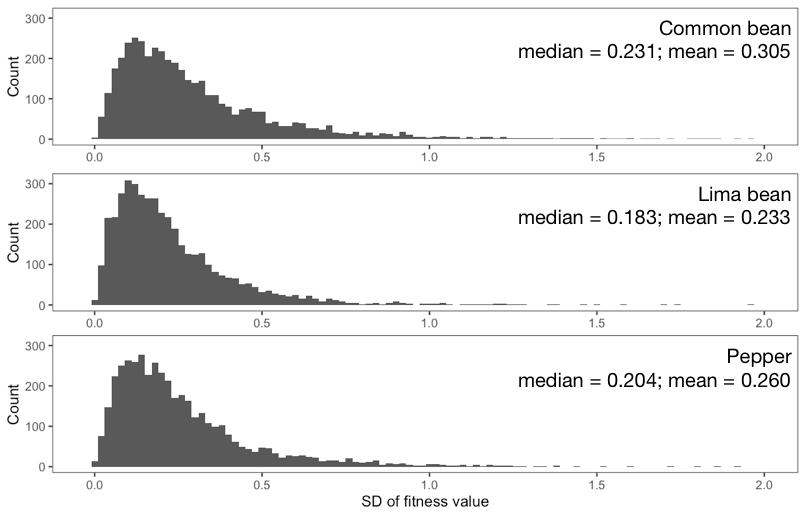

Figure S3. Distribution of standard deviation of gene fitness values for each gene calculated from three replicate *in planta* experiments. These distributions are similar for the three plant species.

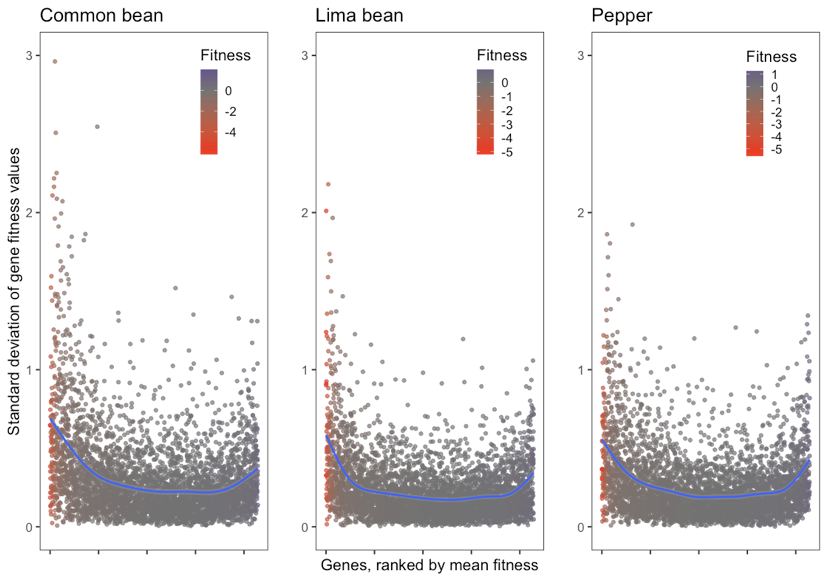

Figure S4. The standard deviation for estimates of gene fitness values is highest at both high and low measures of gene fitness contribution. For each host plant, the standard deviations of gene fitness values are ranked by average gene fitness, and are highest on average for genes with very low or high fitness value. A generalized additive model (GAM) was used to fit the regression lines.

Table S1. Numbers of unique barcodes and median reads per gene obtained from mid-log phase cultures following library outgrowth and before inoculation (time0) and from leaves pooled after growth in 100 pots of plants of a particular host (sample). Unique barcodes were calculated as the total that mapped to the genome and for which 3 or more reads were obtained in a given experiment. The total number of genome-mapped barcodes in the library was 281,417. Technical (sequencing) replicates are listed separately (“a” and “b”), and share the same time0 reference sample. For an experiment to pass quality control, the median reads per gene in the sample must be ≥ 50 [1].

| Experiment | Unique barcodes at time 0 | Unique barcodes in sample | Recovery (%) | Median reads/gene at time 0 | Median reads/gene in sample |
| --- | --- | --- | --- | --- | --- |
| KB_1 | 187,482 | 192,078 | >100* | 278 | 150 |
| KB_2 | 197,089 | 226,857 | >100* | 378.5 | 251 |
| *P.vulgaris*_1a | 218,149 | 149,311 | 68.4 | 409.5 | 163 |
| *P.vulgaris*_1b | 218,149 | 151,211 | 69.3 | 409.5 | 155 |
| *P.vulgaris*_2 | 214,346 | 156,416 | 73.0 | 390 | 160 |
| *P.vulgaris*_3a | 222,473 | 185,183 | 83.2 | 397 | 203 |
| *P.vulgaris*_3b | 222,473 | 187,289 | 84.2 | 397 | 216 |
| *P.lunatus*_1a | 238,593 | 197,334 | 82.7 | 511.5 | 245 |
| *P.lunatus*_1b | 238,593 | 198,378 | 83.1 | 511.5 | 252 |
| *P.lunatus*_2a | 211,661 | 182,104 | 86.0 | 356 | 175 |
| *P.lunatus*_2b | 211,661 | 181,831 | 85.9 | 356 | 166 |
| *P.lunatus*_3a | 208,924 | 199,475 | 95.5 | 368 | 214 |
| *P.lunatus*_3b | 208,924 | 196,817 | 94.2 | 368 | 203 |
| *C.annuum*_1a | 220,024 | 163,572 | 74.3 | 398 | 198 |
| *C.annuum*_1b | 220,024 | 166,370 | 75.6 | 398 | 206 |
| *C.annuum*_2a | 208,449 | 194,666 | 93.4 | 383.5 | 212 |
| *C.annuum*_2b | 208,449 | 201,173 | 96.5 | 383.5 | 220 |
| *C.annuum*_3a | 220,860 | 198,541 | 89.9 | 406 | 219 |
| *C.annuum*_3b | 220,860 | 204,372 | 92.5 | 406 | 239.5 |

* More unique barcodes sequenced at the end of an experiment indicates that additional unique barcodes were present at the threshold abundance in the condition but were below the threshold at the start of the experiment (time0).

Table S2. Unique and shared gene loci among the three hosts tested having average fitness values less than -2 (A) or average fitness values less than -1 (B). These totals are shown as Venn diagrams in Figure 3.

A.

| **Host[s]** | **Total** | **Genes** |
| --- | --- | --- |
| Common bean  Lima bean  Pepper | 34 | Psyr_0033 Psyr_0034 Psyr_0167 Psyr_0219 Psyr_0473 Psyr_0474 Psyr_0529 Psyr_0531 Psyr_0917 Psyr_0918 Psyr_1212 Psyr_1257 Psyr_1269 Psyr_1350 Psyr_1373 Psyr_1613 Psyr_1663 Psyr_1668 Psyr_1669 Psyr_1748 Psyr_1983 Psyr_1984 Psyr_1985 Psyr_2980 Psyr_3008 Psyr_3958 Psyr_4130 Psyr_4270 Psyr_4369 Psyr_4407 Psyr_4580 Psyr_4581 Psyr_4609 Psyr_4991 |
| Common bean  Lima bean | 17 | Psyr_0014 Psyr_0377 Psyr_0378 Psyr_0557 Psyr_0704 Psyr_0826 Psyr_0915 Psyr_1056 Psyr_1408 Psyr_1410 Psyr_1614 Psyr_3179 Psyr_3883 Psyr_4408 Psyr_4683 Psyr_4687 Psyr_4852 |
| Common bean  Pepper | 15 | Psyr_0469 Psyr_0576 Psyr_0846 Psyr_0847 Psyr_0848 Psyr_1217 Psyr_1544 Psyr_2613 Psyr_4132 Psyr_4133 Psyr_4134 Psyr_4893 Psyr_4894 Psyr_4896 Psyr_4897 |
| Lima bean  Pepper | 2 | Psyr_1198 Psyr_4362 |
| Common bean | 17 | Psyr_0454 Psyr_0528 Psyr_0914 Psyr_0919 Psyr_0920 Psyr_0936 Psyr_0951 Psyr_1196 Psyr_2612 Psyr_3637 Psyr_4116 Psyr_4144 Psyr_4340 Psyr_4341 Psyr_4684 Psyr_4686 Psyr_4740 |
| Lima bean | 5 | Psyr_0025 Psyr_0532 Psyr_0923 Psyr_1914 Psyr_4018 |
| Pepper | 7 | Psyr_0385 Psyr_0386 Psyr_1190 Psyr_1191 Psyr_1208 Psyr_1210 Psyr_4566 |

B.

| **Host[s]** | **Total** | **Genes** |
| --- | --- | --- |
| Common bean  Lima bean  Pepper | 90 | Psyr_0014 Psyr_0033 Psyr_0034 Psyr_0167 Psyr_0219 Psyr_0377 Psyr_0378 Psyr_0454 Psyr_0469 Psyr_0473 Psyr_0474 Psyr_0528 Psyr_0529 Psyr_0531 Psyr_0532 Psyr_0557 Psyr_0704 Psyr_0826 Psyr_0827 Psyr_0846 Psyr_0847 Psyr_0848 Psyr_0914 Psyr_0915 Psyr_0917 Psyr_0918 Psyr_0919 Psyr_0920 Psyr_0936 Psyr_0951 Psyr_1056 Psyr_1190 Psyr_1197 Psyr_1198 Psyr_1200 Psyr_1205 Psyr_1206 Psyr_1208 Psyr_1210 Psyr_1211 Psyr_1212 Psyr_1213 Psyr_1216 Psyr_1257 Psyr_1269 Psyr_1350 Psyr_1373 Psyr_1408 Psyr_1410 Psyr_1544 Psyr_1613 Psyr_1663 Psyr_1668 Psyr_1669 Psyr_1747 Psyr_1748 Psyr_1983 Psyr_1984 Psyr_1985 Psyr_2613 Psyr_2980 Psyr_3008 Psyr_3179 Psyr_3636 Psyr_3637 Psyr_3883 Psyr_3958 Psyr_4091 Psyr_4130 Psyr_4132 Psyr_4270 Psyr_4340 Psyr_4341 Psyr_4361 Psyr_4362 Psyr_4369 Psyr_4407 Psyr_4408 Psyr_4566 Psyr_4580 Psyr_4581 Psyr_4609 Psyr_4686 Psyr_4687 Psyr_4852 Psyr_4893 Psyr_4894 Psyr_4896 Psyr_4991 Psyr_5132 |
| Common bean  Lima bean | 25 | Psyr_0025 Psyr_0534 Psyr_0579 Psyr_0916 Psyr_0923 Psyr_1054 Psyr_1055 Psyr_1121 Psyr_1401 Psyr_1614 Psyr_1914 Psyr_2077 Psyr_2461 Psyr_3174 Psyr_3199 Psyr_3287 Psyr_4018 Psyr_4158 Psyr_4194 Psyr_4683 Psyr_4684 Psyr_4843 Psyr_5065 Psyr_5130 Psyr_5133 |
| Common bean  Pepper | 17 | Psyr_0385 Psyr_0386 Psyr_0576 Psyr_1191 Psyr_1196 Psyr_1215 Psyr_1217 Psyr_1218 Psyr_3193 Psyr_4133 Psyr_4134 Psyr_4144 Psyr_4512 Psyr_4844 Psyr_4897 Psyr_4940 Psyr_5072 |
| Lima bean  Pepper | 2 | Psyr_0555 Psyr_1195 |
| Common bean | 57 | Psyr_0103 Psyr_0201 Psyr_0268 Psyr_0475 Psyr_0533 Psyr_0550 Psyr_0758 Psyr_0831 Psyr_1097 Psyr_1109 Psyr_1247 Psyr_1395 Psyr_1417 Psyr_1419 Psyr_1487 Psyr_1542 Psyr_1588 Psyr_1733 Psyr_1751 Psyr_1907 Psyr_1998 Psyr_2221 Psyr_2264 Psyr_2396 Psyr_2474 Psyr_2501 Psyr_2557 Psyr_2601 Psyr_2612 Psyr_3028 Psyr_3427 Psyr_3552 Psyr_3597 Psyr_3667 Psyr_3675 Psyr_3676 Psyr_3678 Psyr_3690 Psyr_3698 Psyr_3791 Psyr_3889 Psyr_4015 Psyr_4044 Psyr_4069 Psyr_4116 Psyr_4125 Psyr_4136 Psyr_4143 Psyr_4224 Psyr_4740 Psyr_4754 Psyr_4774 Psyr_4882 Psyr_4895 Psyr_5053 Psyr_5067 Psyr_5129 |
| Lima bean | 28 | Psyr_0202 Psyr_0259 Psyr_0435 Psyr_0478 Psyr_0524 Psyr_0796 Psyr_0822 Psyr_1053 Psyr_1057 Psyr_1058 Psyr_1059 Psyr_1060 Psyr_1061 Psyr_1062 Psyr_1063 Psyr_1140 Psyr_1667 Psyr_1749 Psyr_2462 Psyr_3146 Psyr_3684 Psyr_3691 Psyr_4008 Psyr_4019 Psyr_4100 Psyr_4627 Psyr_4842 Psyr_4898 |
| Pepper | 8 | Psyr_1111 Psyr_2080 Psyr_2245 Psyr_2543 Psyr_2617 Psyr_3459 Psyr_3835 Psyr_5135 |

Table S3. Genes having a differential fitness contribution to growth in three plant species by a Kruskal-Wallis rank sum test (p < 0.05). Fitness values in KB are included for comparison.

| Locus | Name | Description | Classification | Average fitness | | | |
| --- | --- | --- | --- | --- | --- | --- | --- |
|  |  |  |  | KB | Common bean | Lima bean | Pepper |
| Psyr_0025 | aroE | shikimate dehydrogenase | Amino acid metabolism and transport | -0.25 | -1.62 | -2.95 | -0.31 |
| Psyr_4133 | hisD | histidinol dehydrogenase | Amino acid metabolism and transport | 0.12 | -2.99 | -0.83 | -3.80 |
| Psyr_4896 | hisH | imidazole glycerol phosphate synthase subunit hisH | Amino acid metabolism and transport | -0.10 | -2.20 | -1.06 | -3.23 |
| Psyr_4852 |  | D-3-phosphoglycerate dehydrogenase | Amino acid metabolism and transport | -0.13 | -2.94 | -3.40 | -1.95 |
| Psyr_0758 | scrB | beta-fructofuranosidase | Carbohydrate metabolism and transport | 0.05 | -1.66 | -0.94 | -0.52 |
| Psyr_2188 |  | Histidine kinase, HAMP region:Bacterial chemotaxis sensory transducer:CHASE3 | Chemosensing & chemotaxis | -0.23 | -0.88 | 0.18 | -0.15 |
| Psyr_4687 | bioB | biotin synthase | Cofactor metabolism | -2.94 | -2.86 | -2.24 | -1.63 |
| Psyr_4684 | bioC | biotin synthesis protein BioC | Cofactor metabolism | -2.76 | -2.85 | -1.79 | -0.87 |
| Psyr_4686 | bioF | 8-amino-7-oxononanoate synthase | Cofactor metabolism | -3.03 | -2.62 | -1.99 | -1.30 |
| Psyr_2098 | cobA | uroporphyrinogen-III C-methyltransferase | Cofactor metabolism | 0.27 | -0.56 | 0.08 | -0.23 |
| Psyr_0847 | ilvH | acetolactate synthase, small subunit | Cofactor metabolism | 0.08 | -2.45 | -1.44 | -3.89 |
| Psyr_0846 | ilvI | acetolactate synthase, large subunit | Cofactor metabolism | -0.18 | -3.10 | -1.92 | -3.70 |
| Psyr_3487 | flgA | flagellar protein FlgA | Flagellar synthesis and motility | 0.14 | 0.27 | 0.02 | 0.78 |
| Psyr_0487 | gshB | glutathione synthase | Glutathione metabolism | -1.54 | 1.81 | 0.78 | -0.34 |
| Psyr_1716 |  | conserved hypothetical protein | Hypothetical | 0.26 | -0.87 | -0.40 | 0.00 |
| Psyr_2543 |  | conserved hypothetical protein | Hypothetical | 0.57 | -0.07 | 0.34 | -1.05 |
| Psyr_3107 |  | conserved hypothetical protein | Hypothetical | -0.18 | -0.54 | 0.20 | 0.44 |
| Psyr_3172 |  | Glycosyl transferase, family 3 | Hypothetical | -0.04 | 0.52 | -0.30 | 0.11 |
| Psyr_3611 |  | Protein of unknown function DUF815 | Hypothetical | 0.22 | 0.68 | -0.56 | 0.10 |
| Psyr_3889 |  | conserved hypothetical protein | Hypothetical | -0.01 | -1.39 | -0.06 | 0.52 |
| Psyr_0014 |  | lipid A biosynthesis acyltransferase | LPS synthesis and transport | -0.53 | -2.66 | -2.08 | -1.44 |
| Psyr_3369 |  | Twin-arginine translocation pathway signal:Tat-translocated enzyme:Dyp-type peroxidase | Oxidative stress tolerance (Antioxidant enzyme) | -0.10 | -0.55 | 0.22 | -0.25 |
| Psyr_2621 | pseB | Secretion protein HlyD | Phytotoxin synthesis and transport | -0.06 | 0.25 | -0.03 | -0.53 |
| Psyr_2601 | salA | regulatory protein, LuxR | Phytotoxin synthesis and transport | -0.37 | -1.21 | -0.23 | -0.68 |
| Psyr_1702 | sylA | regulatory protein, LuxR | Phytotoxin synthesis and transport | -0.26 | -0.54 | -0.10 | 0.18 |
| Psyr_2614 | sypA | Amino acid adenylation | Phytotoxin synthesis and transport | 0.03 | -0.85 | -0.15 | -0.69 |
| Psyr_2612 | syrP | syrP protein, putative | Phytotoxin synthesis and transport | 0.04 | -2.07 | -0.02 | -0.70 |
| Psyr_4158 | eftA | conserved hypothetical protein | Plant-associated proteins | -0.39 | -1.42 | -1.01 | -0.09 |
| Psyr_1061 | alg44 | alginate biosynthesis protein Alg44 | Polysaccharide synthesis and regulation | 0.02 | -0.72 | -1.28 | -0.34 |
| Psyr_1062 | alg8 | alginate biosynthesis protein Alg8 | Polysaccharide synthesis and regulation | -0.04 | -0.70 | -1.22 | -0.33 |
| Psyr_0937 | algA-1 | mannose-6-phosphate isomerase, type 2 / mannose-1-phosphate guanylyltransferase (GDP) | Polysaccharide synthesis and regulation | -0.30 | -0.70 | -0.47 | -0.16 |
| Psyr_1053 | algF | alginate biosynthesis protein AlgF | Polysaccharide synthesis and regulation | -0.04 | -0.81 | -1.48 | -0.43 |
| Psyr_1055 | algI | Membrane bound O-acyl transferase, MBOAT | Polysaccharide synthesis and regulation | 0.00 | -1.03 | -1.57 | -0.36 |
| Psyr_1060 | algK | Sel1-like repeat protein | Polysaccharide synthesis and regulation | 0.17 | -0.51 | -1.21 | -0.28 |
| Psyr_0378 | mdoH | Glycosyl transferase, family 2 | Polysaccharide synthesis and regulation | -1.04 | -3.80 | -4.04 | -1.08 |
| Psyr_3636 | wbpM | Polysaccharide biosynthesis protein CapD | Polysaccharide synthesis and regulation | -0.45 | -1.76 | -1.48 | -1.10 |
| Psyr_3161 | aprD | Type I secretion system ATPase, PrtD | Secretion/Efflux/Export | 0.39 | 0.11 | 0.38 | -0.61 |
| Psyr_4009 | oprM | RND efflux system, outer membrane lipoprotein, NodT | Secretion/Efflux/Export | 0.02 | -0.52 | -0.95 | 0.20 |
| Psyr_0831 | cbrB-1 | Two-component response regulator CbrB | Signal transduction mechanisms | -1.45 | -1.71 | -0.98 | 0.17 |
| Psyr_4069 | colS | ATP-binding region, ATPase-like:Histidine kinase, HAMP region:Histidine kinase A, N-terminal | Signal transduction mechanisms | -0.08 | -1.21 | -0.57 | -0.38 |
| Psyr_4138 |  | Toluene tolerance | Stress resistance | -0.22 | -0.20 | -0.59 | 0.28 |
| Psyr_3698 | gacS | Response regulator receiver:ATP-binding region, ATPase-like:Histidine kinase, HAMP region:Histidine kinase A, N-terminal:Hpt | Transcriptional regulation | 0.07 | -1.47 | -0.44 | -0.85 |
| Psyr_4239 | dppB | Binding-protein-dependent transport systems inner membrane component | Transport (peptides) | -0.09 | -0.58 | -0.33 | -0.07 |
| Psyr_4240 | dppC | Binding-protein-dependent transport systems inner membrane component | Transport (peptides) | 0.14 | -0.57 | -0.34 | 0.01 |
| Psyr_1218 | hrpK1 | type III helper protein HrpK1 | Type III secretion system | 0.08 | -1.28 | -0.75 | -1.67 |
| Psyr_0914 | wpbZ | Glycosyl transferase, group 1 |  | -0.19 | -2.08 | -1.83 | -1.32 |
| Psyr_0915 |  | NAD-dependent epimerase/dehydratase |  | -0.21 | -3.19 | -3.51 | -1.36 |
| Psyr_1419 |  | preQ(0) biosynthesis protein QueC |  | -0.65 | -1.85 | -0.87 | -0.10 |
| Psyr_4844 |  | HAD-superfamily hydrolase, subfamily IB (PSPase-like):HAD-superfamily subfamily IB hydrolase, hypothetical 2 |  | -0.10 | -1.00 | -0.30 | -1.69 |
| Psyr_4886 |  | Peptidase M23B |  | -0.05 | -0.53 | -0.30 | -0.07 |

This table does not include 37 genes whose average fitness values differed between plant species but whose effect was small, having fitness values greater than -0.5 and less than +0.5 in all host plant species.

Table S4. Genes having a differential fitness contribution to growth in three plant species by a Kruskal-Wallis rank sum test (p < 0.05), but whose effect was small, having fitness values greater than -0.5 and less than +0.5 in all host plant species. Fitness values in KB are included for comparison.

| Locus | Name | Description | Classification | Average fitness | | | |
| --- | --- | --- | --- | --- | --- | --- | --- |
|  |  |  |  | KB | Common bean | Lima bean | Pepper |
| Psyr_1074 | aapM | amino acid ABC transporter membrane protein 2, PAAT family | Amino acid metabolism and transport | 0.17 | 0.10 | 0.28 | -0.06 |
| Psyr_3876 | hisM | amino acid ABC transporter membrane protein 2, PAAT family | Amino acid metabolism and transport | -0.04 | 0.23 | 0.08 | -0.09 |
| Psyr_2715 |  | Major facilitator superfamily | Carbohydrate metabolism and transport | 0.02 | 0.30 | -0.16 | 0.12 |
| Psyr_3406 | aer-2 | PAS | Chemosensing & chemotaxis | -0.06 | -0.29 | 0.05 | -0.13 |
| Psyr_2995 | treY | maltooligosyl trehalose synthase | Compatible solute synthesis | 0.02 | -0.45 | -0.18 | 0.01 |
| Psyr_1481 | ppa-2 | Inorganic diphosphatase | Energy generation | -0.09 | -0.06 | 0.03 | -0.20 |
| Psyr_1770 |  | Enoyl-CoA hydratase/isomerase | Fatty acid metabolism | -0.07 | -0.06 | -0.27 | 0.30 |
| Psyr_3456 | fliG | Flagellar motor switch protein FliG | Flagellar synthesis and motility | 0.09 | -0.24 | 0.04 | 0.38 |
| Psyr_3613 |  | Glutathione peroxidase | Glutathione metabolism | 0.12 | 0.39 | -0.19 | 0.08 |
| Psyr_0332 |  | hypothetical protein | Hypothetical | -0.06 | 0.21 | -0.10 | 0.05 |
| Psyr_1137 |  | Protein of unknown function UPF0153 | Hypothetical | 0.00 | -0.12 | 0.26 | 0.12 |
| Psyr_1407 |  | Protein of unknown function DUF28 | Hypothetical | -0.07 | 0.41 | 0.07 | -0.31 |
| Psyr_1533 |  | hypothetical protein | Hypothetical | 0.16 | -0.39 | -0.11 | 0.15 |
| Psyr_2339 |  | hypothetical protein | Hypothetical | 0.09 | 0.46 | -0.37 | 0.18 |
| Psyr_2947 |  | hypothetical protein | Hypothetical | -0.22 | 0.47 | -0.01 | -0.39 |
| Psyr_3006 |  | Protein of unknown function DUF419 | Hypothetical | -0.10 | 0.40 | -0.10 | 0.06 |
| Psyr_3066 |  | conserved hypothetical protein | Hypothetical | -0.22 | -0.10 | -0.39 | 0.27 |
| Psyr_3740 |  | Protein of unknown function DUF454 | Hypothetical | 0.03 | -0.33 | 0.01 | 0.13 |
| Psyr_3798 |  | conserved domain protein | Hypothetical | 0.01 | 0.16 | 0.01 | -0.09 |
| Psyr_4248 |  | hypothetical protein | Hypothetical | 0.01 | 0.08 | 0.14 | -0.01 |
| Psyr_5112 |  | conserved hypothetical protein | Hypothetical | -0.13 | 0.03 | -0.06 | 0.16 |
| Psyr_1318 | ppc | Phosphoenolpyruvate carboxylase | Organic acid metabolism and transport | 0.15 | -0.18 | 0.22 | 0.09 |
| Psyr_1705 | sylD | Amino acid adenylation | Phytotoxin synthesis and transport | 0.03 | -0.32 | -0.02 | -0.06 |
| Psyr_4615 |  | Spermidine/putrescine ABC transporter ATP-binding subunit | Polyamine metabolism and transport | -0.09 | -0.15 | 0.10 | 0.20 |
| Psyr_3956 | mucB | sigma E regulatory protein, MucB/RseB | Polysaccharide synthesis and regulation | -0.04 | 0.06 | 0.01 | 0.33 |
| Psyr_3236 | dhcR | transcriptional regulator, LysR family | QAC metabolism and transport | 0.32 | -0.12 | 0.06 | 0.29 |
| Psyr_2576 | syfA | Amino acid adenylation | Secondary metabolism | 0.10 | -0.03 | 0.10 | 0.31 |
| Psyr_2864 |  | RND efflux system, outer membrane lipoprotein, NodT | Secretion/Efflux/Export | 0.14 | -0.11 | -0.01 | -0.28 |
| Psyr_3989 | xaxA | hypothetical protein | Special | -0.02 | 0.15 | 0.08 | -0.02 |
| Psyr_4273 | cstA | Carbon starvation protein CstA | Stress resistance | 0.05 | -0.05 | 0.04 | 0.15 |
| Psyr_0986 | rsmC | 16S rRNA m(2)G 1207 methyltransferase |  | -0.19 | -0.37 | 0.00 | 0.37 |
| Psyr_0929 |  | Glycosyl transferase, family 2 |  | 0.11 | -0.20 | -0.07 | -0.02 |
| Psyr_0989 |  | Lysine exporter protein (LYSE/YGGA) |  | 0.06 | -0.25 | -0.03 | 0.32 |
| Psyr_3859 |  | Purine nucleoside permease |  | -0.01 | -0.23 | -0.08 | 0.04 |
| Psyr_4631 |  | PrkA serine kinase |  | 0.07 | -0.09 | -0.03 | 0.07 |
| Psyr_5082 |  | Band 7 protein |  | -0.02 | -0.01 | -0.24 | 0.11 |
| Psyr_5111 |  | dTDP-glucose 4,6-dehydratase |  | -0.06 | 0.08 | -0.03 | -0.13 |

Table S5. Strains used in this study.

| Strains | Genotype | Reference |
| --- | --- | --- |
| *P. syringae* B728a | Wild type strain (Rif^R^) | [2] |
| *P. syringae* B728a | Whole genome barcoded *mariner* transposon library (Rif^R^ Kan^R^) | [3] |

**Literature Cited**

1.	Wetmore KM, Price MN, Waters RJ, Lamson JS, He J, Hoover CA, et al. Rapid quantification of mutant fitness in diverse bacteria by sequencing randomly bar-coded transposons. MBio. 2015;6: 1–15. doi:10.1128/mBio.00306-15

2.	Loper JE, Lindow SE. Lack of evidence for *in situ* fluorescent pigment production by *Pseudomonas syringae* pv. *syringae* on bean leaf surfaces. Phytopathology. 1987;77: 1449–1454. doi:10.1094/Phyto-77-1449

3.	Helmann TC, Deutschbauer AM, Lindow SE. Genome-wide identification of *Pseudomonas syringae* genes required for fitness during colonization of the leaf surface and apoplast. Proc Natl Acad Sci. 2019;116: 18900–18910. doi:10.1073/pnas.1908858116
